# Discovering Novel Prognostic Biomarkers of Hepatocellular Carcinoma using eXplainable Artificial Intelligence

**DOI:** 10.1101/2023.11.27.568859

**Authors:** Elizabeth Gutierrez-Chakraborty, Debaditya Chakraborty, Debodipta Das, Yidong Bai

## Abstract

Hepatocellular carcinoma (HCC) remains a global health challenge with high mortality rates, largely due to late diagnosis and suboptimal efficacy of current therapies. With the imperative need for more reliable, non-invasive diagnostic tools and novel therapeutic strategies, this study focuses on the discovery and application of novel genetic biomarkers for HCC using explainable artificial intelligence (XAI). Despite advances in HCC research, current biomarkers like Alpha-fetoprotein (AFP) exhibit limitations in sensitivity and specificity, necessitating a shift towards more precise and reliable markers. This paper presents an innovative XAI framework to identify and validate key genetic biomarkers for HCC prognosis. Our methodology involved analyzing clinical and gene expression data to identify potential biomarkers with prognostic significance. The study utilized robust AI models validated against extensive gene expression datasets, demonstrating not only the predictive accuracy but also the clinical relevance of the identified biomarkers through explainable metrics. The findings highlight the importance of biomarkers such as TOP3B, SSBP3, and COX7A2L, which were consistently influential across multiple models, suggesting their role in improving the predictive accuracy for HCC prognosis beyond AFP. Notably, the study also emphasizes the relevance of these biomarkers to the Hispanic population, aligning with the larger goal of demographic-specific research. The application of XAI in biomarker discovery represents a significant advancement in HCC research, offering a more nuanced understanding of the disease and laying the groundwork for improved diagnostic and therapeutic strategies.

## 1. Introduction

Hepatocellular carcinoma (HCC) is a leading cause of cancer-related mortality and is currently the main cause of liver-related death, leading to more than a million deaths annually worldwide (*1*–*3*). Over several decades, substantial progress has been made in the understanding of HCC risk factors, epidemiology, and molecular pathogenesis. However, HCC still poses a significant health challenge due to its typically late presentation and the limited success of current therapies (*4*–*7*). The late onset of symptoms, often during advanced disease stages, complicates timely diagnosis and effective intervention. Consequently, the development of accessible, non-invasive diagnostic methods and innovative treatments is critical to improving patient care and outcomes in HCC (*8*). There is an urgent need for easy diagnostic and prognostic tools, and novel therapies for the management of HCC that would greatly improve patient management and clinical outcomes. Biomarkers are pivotal in the HCC diagnostic process, assisting in early detection, prognosis assessment, treatment monitoring, and post-treatment recurrence surveillance. Alpha-fetoprotein (AFP) is an established biomarker for HCC (*9*) even though it has been reported that up to 20-40% of HCC patients’ tumor cells do not secrete AFP proteins (*10*–*12*). Furthermore, elevated AFP levels are not exclusive to HCC and may arise from other hepatic pathologies or germ cell tumors. Research indicates that the application of AFP is limited due to its low sensitivity and specificity (*13*), and approximately 15-30% of advanced HCC cases do not exhibit elevated AFP levels (*14*). As such, reliance on AFP as a solitary diagnostic tool is diminishing due to its low sensitivity and specificity thus limiting its clinical use (*15, 16*). Therefore, AFP testing does not have much of a role in the diagnosis of HCC (*17*). Reflecting this, the American Association for the Study of Liver Diseases (AASLD) has revised its guidelines, advising against the exclusive use of AFP for early HCC detection (*14*), underscoring the necessity for more reliable biomarkers. Additionally, a meta-analysis of cohort studies reported the sensitivity of ultrasound for early-stage HCC detection is only 45%, which increases to 63% with the addition of AFP (*7, 18*). Despite some advancements in diagnosing and managing HCC, its overall prognosis continues to be poor (*19*). This ongoing challenge highlights the critical need for developing dependable biomarkers not only vital for improving diagnostic accuracy and treatment strategies but also crucial for the effective prognostic assessment of HCC (*20*).

Considering the overall survival rate is alarmingly low at 10-20% (*21*), identifying new biomarkers for early and precise detection, prognosis, and therapeutic targeting across all HCC variants is of paramount importance (*22*). The potential for these biomarkers to serve as noninvasive diagnostics or the basis for drug development is significant. Despite a decade of advances in cancer research, patient outcomes have seen little improvement (*23*), with HCC accounting for approximately 90% of primary liver cancer cases (*24*). HCC also exhibits a higher incidence in Hispanic and Asian/Pacific Islander populations compared to non-Hispanic whites, indicating a disproportionate impact on these groups (*25*). The current challenge is the unsatisfactory survival rate for HCC patients (*3*), prompting ongoing research into factors influencing HCC survival, tumor progression, and metastasis. New findings in these areas could lead to novel and more effective treatments. While groundbreaking scientific insights have been achieved, translation into clinical practice is often a slow and costly process. Traditional research methods may be prone to design bias, may disproportionately focus on certain molecules, and risk missing more effective biomarkers (*26*). However, the application of eXplainable Artificial Intelligence (XAI) provides a promising alternative to traditional methods. XAI offers a way to expedite the discovery process, reduce costs, and mitigate research biases, enabling the identification of valuable insights into HCC with greater efficiency and potentially transforming the landscape of liver cancer treatment.

Employing a suite of tree-based ensemble Artificial Intelligence (AI) models, including Extreme Gradient Boosting (XGBoost), Random Forest (RFC), and Extra Trees Classifiers (ETC), coupled with SHAP for post-hoc explainability, enabled the identification of key genetic biomarkers with a significant impact on HCC prognosis. The integration of findings across these models pinpointed a set of critical genes influencing five-year survival rates in HCC patients. This study highlights SSBP3 as a consistently influential (one of the top five) gene across all AI models utilized, indicating its potential as a critical biomarker in HCC prognosis. Additionally, COX7A2L has been identified as influential by two of the three AI models, further underscoring its possible significance in the disease’s progression. These findings underscore that a composite application of these AI-identified biomarkers markedly enhanced prognostic accuracy beyond the capabilities of existing markers, such as AFP, currently utilized in HCC detection.

## 2. Materials and Methods

**Figure 1** illustrates the analytical framework employed in our study to discern and corroborate genetic biomarkers for HCC. The methodology commences with the retrieval of clinical and gene expression data from public repositories. Subsequently, this data is processed through the lens of XAI, enabling the identification of novel HCC biomarkers that can potentially serve as indicators of survival outcomes and therapeutic targets. These biomarkers are then represented through a series of graphs, elucidating patterns such as survival rates correlated with biomarker levels and gene expression distributions. Moreover, the study incorporates a localized dataset (*27*), encompassing both Hispanic and general population profiles. This local data is analyzed to pinpoint genes of particular interest, especially those relevant to the Hispanic population, which might have a different prevalence rate or genetic expression pattern of HCC.

**Figure 1.**
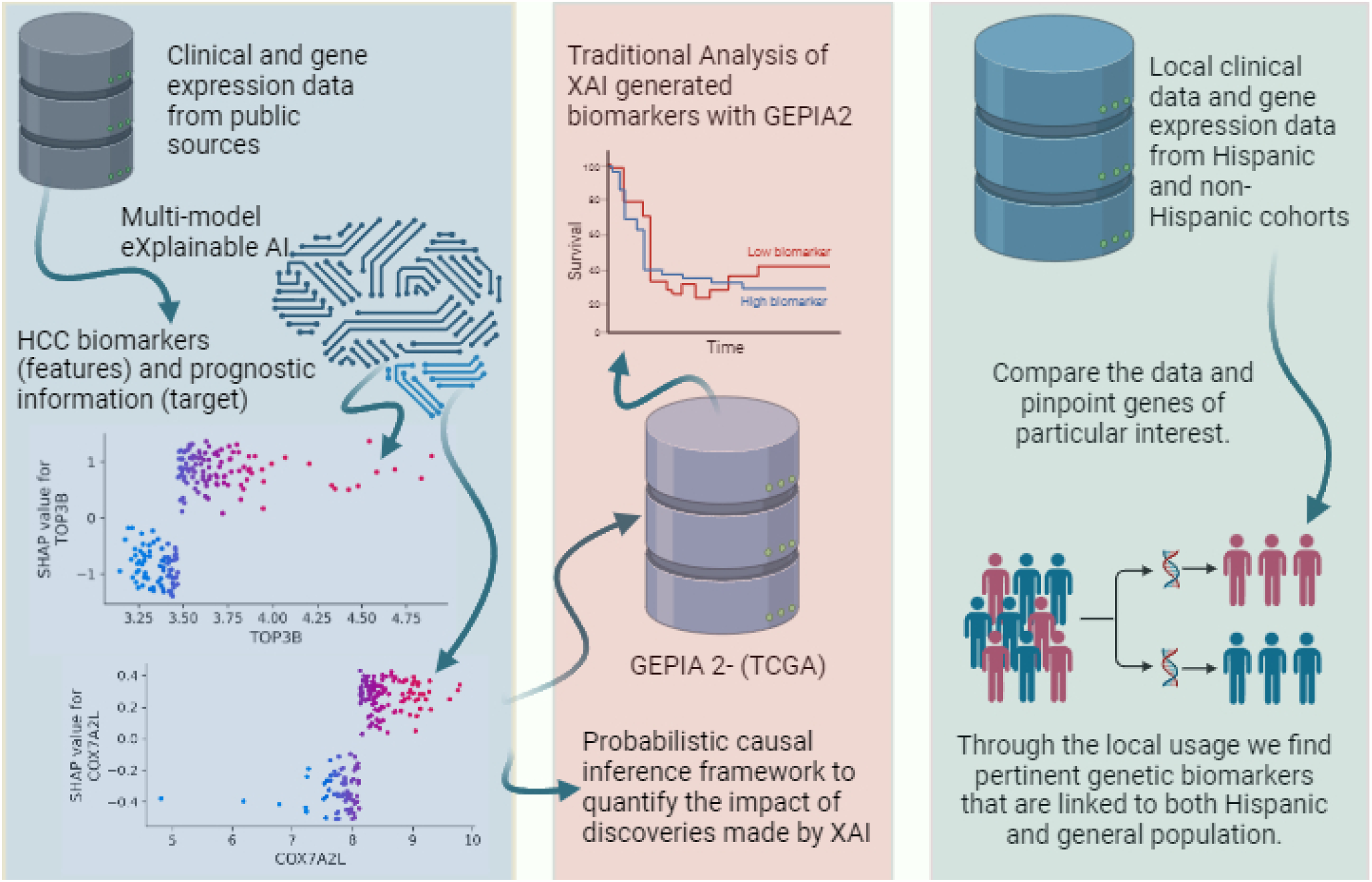
Presents a schematic overview of the research methodology. Utilizing clinical and gene expression data sourced from public repositories, the study primarily employs XAI to discover HCC biomarkers.

Our study harnesses an XAI framework to forecast survival and clinical outcomes in HCC patients, analyzing genetic data through predictive AI modeling. This approach facilitates the detection and graphical representation of pivotal biomarkers, while also explaining their biological relevance in HCC progression. We employed the comprehensive HCCDB: A Database of Hepatocellular Carcinoma Expression Atlas (*28*), which encompasses 22 normal liver tissue samples and 231 total samples featuring 12,571 genes, to develop, fine-tune, and evaluate our innovative XAI models. The database has curated RNA-Seq and microarray-based data from multiple sources, including TCGA and GEO. The models were swiftly trained and validated, with XGBoost requiring just 0.03 minutes and RFC and ETC each taking 0.01 minutes, including data loading and model refinement through five-fold cross-validation. The XAI-generated insights distinctly rank the biomarkers in an interpretable manner, spotlighting the most significant genes for prognostic assessment in HCC patient care.

### Knowledge discovery with XAI

The XAI model was developed by combining inherently interpretable tree-based ensembled AI algorithms (*29*), with a post hoc game theory-based explainer model known as SHAP (*30*), which have recently been shown to be successful on structured datasets in multiple biological domains (*31*–*33*).

Conceptually, the ensembled algorithms that are developed by boosting individual trees learn the functional relationship (*f*) between the features (*X*) and target (*Y*) through an iterative process in which the individual trees are sequentially trained on the residuals from the previous tree. Mathematically, the predictions from the trees can be expressed as:

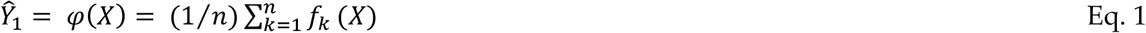

In Eq. 1, 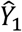 are the predictions, 1 ≤ *k* ≤ *n*, and *n* is the total number of functions learned by the *n* number of trees. The regularized objective *L*(φ) is minimized to learn the set of functions *f*_*k*_ used in the model.

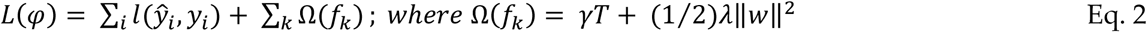

In Eq. 2, *l* is a differentiable convex loss function that measures the difference between 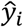 (prediction) and *y*_*i*_ (target). Ω is an extra regularization term that penalizes the growing of more trees in the model to prevent complexity and thus, reduce overfitting. *γ* is the complexity of each leaf, *T* is the number of leaves in a tree, λ is a penalty parameter, and ‖*w*‖ is the vector of scores on the leaves.

Additionally, we applied a bagging-based ensembling technique in which each tree is trained with values of a random vector sampled independently and with the same distribution for all other trees in the forest. From a conceptual point of view, the *k*^*th*^ tree is trained using a random vector Θ_*k*_ that is independent of the past random vectors Θ_1_ to Θ_*k*−1_ but with the same distribution, resulting in a tree *h*(*X*, Θ_*k*_) where *X* is an input vector. When a large number of trees are grown in the ensemble, their average predictions will be taken, which will improve the predictive accuracy and control over-fitting. Mathematically, this is represented as:

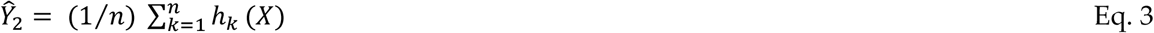

In Eq. 3, 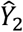 are the predictions, in this case, 1 ≤ *k* ≤ *n*, and *n* is the total number of trees generated. The mean-squared generalization error of any tree *h*(*X*) is given by *E*_*X,Y*_(*Y* − *h*(*X*))^2^ for the input vector *X* and the target *Y*. As the number of trees *h*(*X*) in the ensemble goes to infinity, *E*_*X,Y*_(*Y* − *av*_*k*_*h*(*X*, Θ_*k*_))^2^ tends to *E*_*X,Y*_(*Y* − *E*_Θ_*h*(*X*, Θ))^2^. This approach generates training subsets in which bootstrap samples are drawn with replacement. Finally, the predictions produced by the ensembled models explained above will be aggregated to produce the final predictions 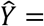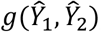, where *g* is a function to aggregate the prediction probabilities and assign classes corresponding to the highest probability. The interpretability and explainability of ensemble models are enhanced through post-hoc feature attribution techniques, employing a threefold approach: (i) employing a polynomial-time algorithm rooted in game theory for calculating precise explanations, (ii) using explanations that quantify the effect of local feature interactions, and (iii) utilizing tools that aggregate numerous local explanations to illuminate the model’s global structure (*30*). Global explanations yield an overarching insight into feature impact on model predictions, whereas local explanations delve into the nuanced ways predictions change in response to specific feature values and their interplay.

### Statistical metrics to report the models’ predictive performances

We have utilized four standard statistical metrics for classification tasks that are mathematically defined below:

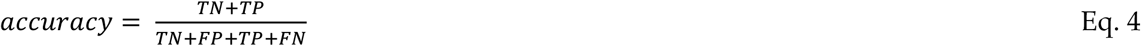

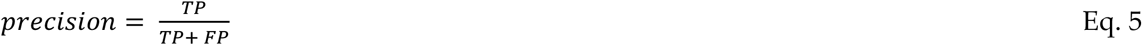

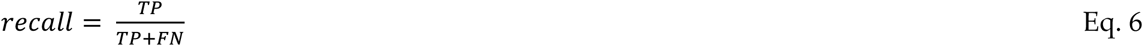

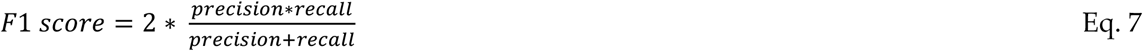

In this analysis, ‘true negatives’ (TN) denote the count of patients correctly identified as surviving beyond five years, while ‘true positives’ (TP) reflect the accurate classification of patients who survive less than five years. Conversely, ‘false negatives’ (FN) are those patients incorrectly predicted to survive beyond five years, and ‘false positives’ (FP) are the instances where patients are wrongly classified as not reaching the five-year survival threshold. In our coding system, survival beyond five years is encoded as ‘0’, and survival under five years as ‘1’, which aligns with the representation in the confusion matrix in **Figure 2**. This figure also compares the performance of three modeling techniques: XGBoost, RFC, and ETC. The results indicate that the bagging methods, like Random Forest and Extra Trees, achieve fewer misclassifications than the boosting method employed by XGBoost.

**Figure 2.**
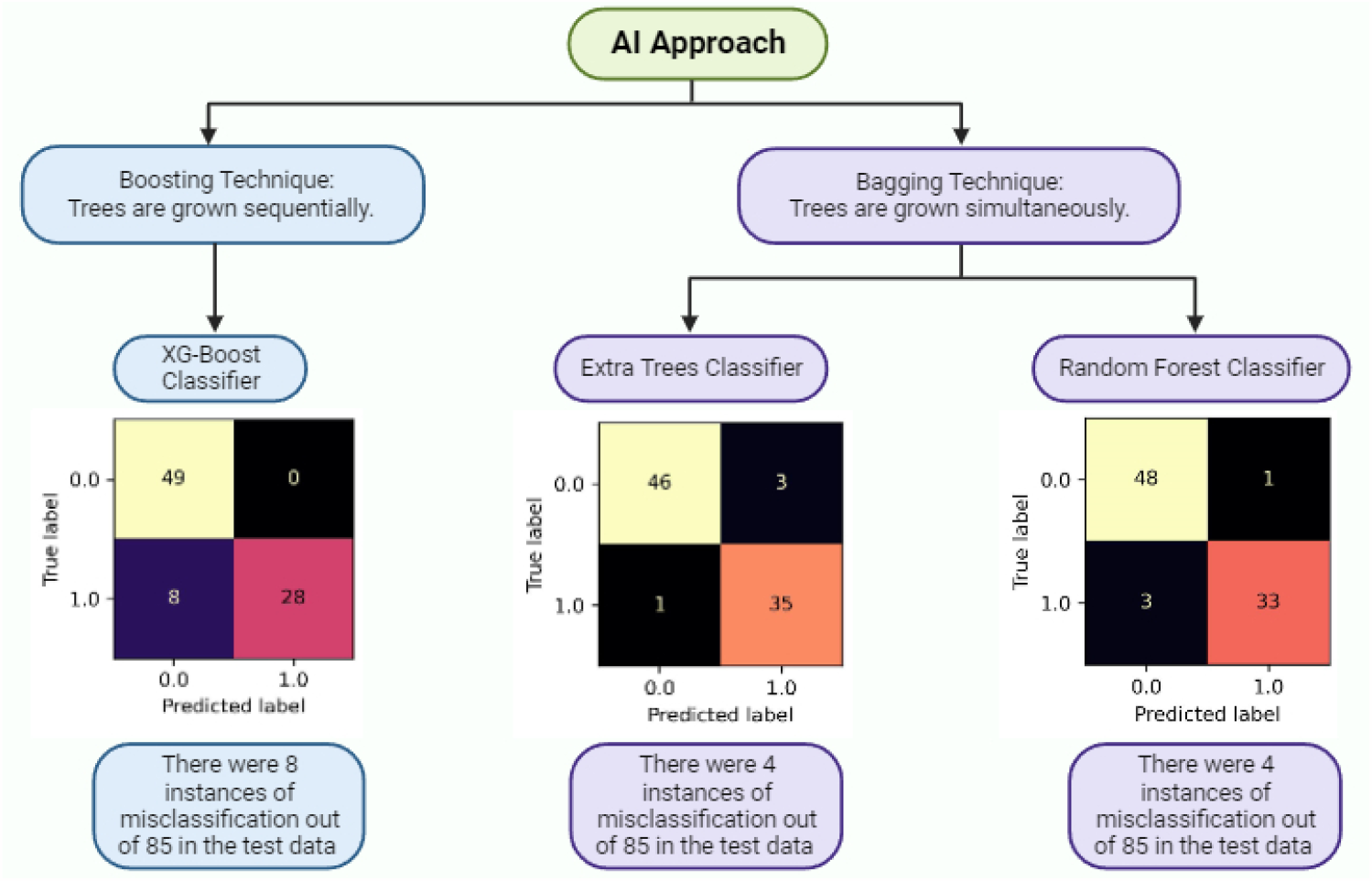
This figure illustrates the comparative performance of the three distinct AI algorithms employed in this study. The algorithm that exhibits the lowest rate of misclassifications has better predictability relative to the others. This visualization aids in discerning the most predictive AI approach for prognostic modeling in our research.

## 3. Results

### Insights generated by XAI

In the previous section, we discussed how integrating predictive AI models with post-hoc feature attribution methods offers detailed insights into the genetic determinants of a patient’s likelihood of surviving HCC for five years or more. XAI-generated global explanations delineate the relationship between gene expression variability and patient prognoses. **Figure 3** A, B, and C rank gene importance in descending order, with the most influential genes for five-year survival at the top emerging from each of the AI models utilized. Positive values on the x-axis suggest an increased risk of mortality within five years, while negative values are indicative of a favorable five-year survival prognosis. Color coding further enhances interpretability, with red denoting higher gene expression and blue indicating lower. For instance, SSBP3’s high expression—associated with favorable outcomes—is consistently marked in red and located on the left, suggesting its role as a possible cancer suppressor. Conversely, genes expressions marked in red that appear on the right, such as TOP3B and COX7A2L, may be implicated as oncogenic or cancer promoter. Such data demonstrate the potential of these genes to serve as diagnostic and prognostic markers, and therapeutic targets.

**Figure 3.**
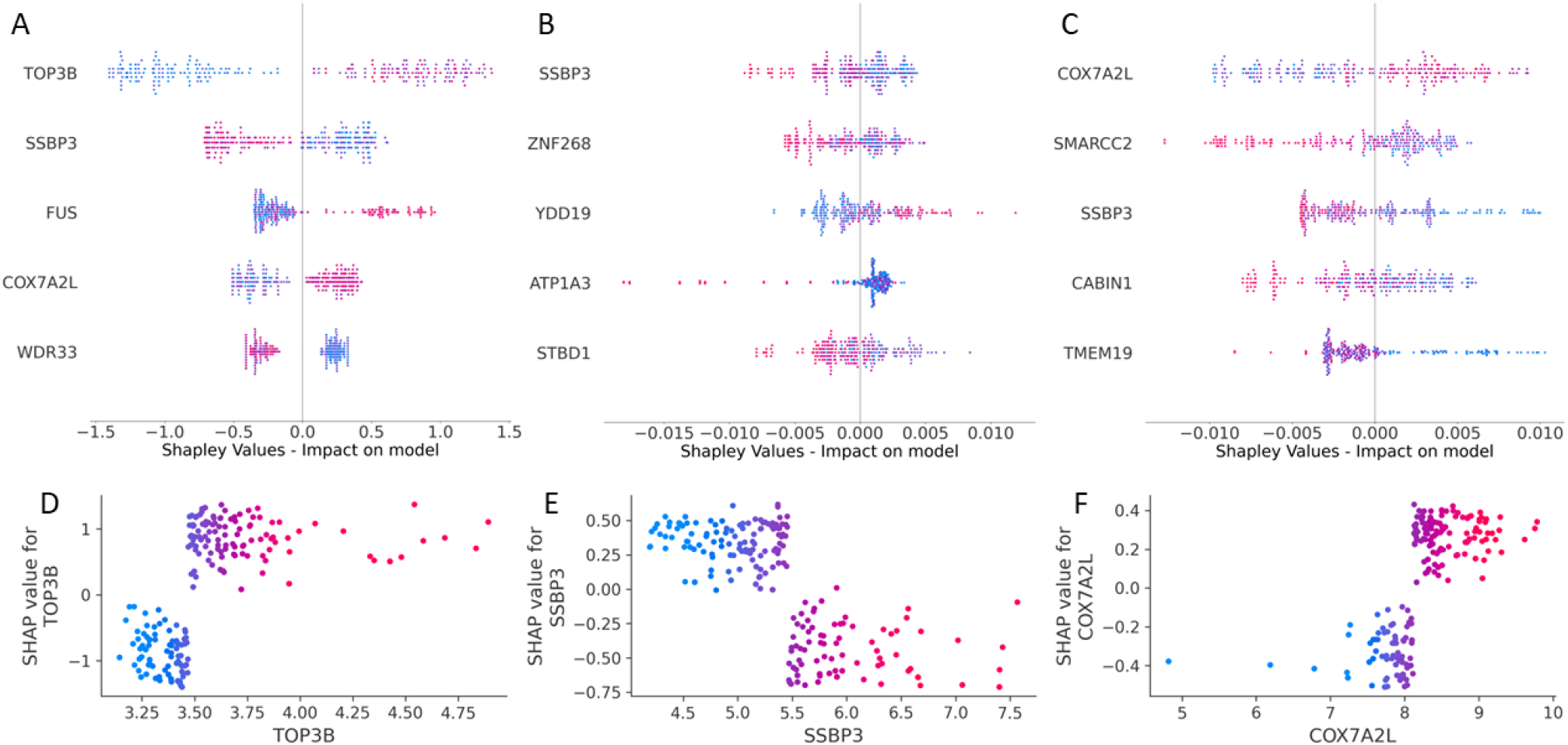
Panels **A, B**, and **C** depicts the global explanatory power of XGBoost, ETC, and RFC models respectively, utilizing SHAP values to quantify the impact of gene expression variability on the prognosis of HCC patients. These global insights aggregate the influence of multiple genes, such as TOP3B, SSBP3, and COX7A2L, on model outcomes. Panels **D, E**, and **F** provide focused local explanations, pinpointing the inflection points for the respective genes TOP3B, SSBP3, and COX7A2L. These local explanations illustrate the specific gene expression thresholds that significantly sway the prognostic predictions. These local plots demarcate how gene expressions contribute to prognoses, depicted by red and blue dots to denote higher and lower gene expressions, respectively. The SHAP/Shapley values are mapped to positive and negative axes, corresponding to less favorable and more favorable patient outcomes, respectively, providing a nuanced understanding of gene impacts on HCC survival predictions.

Employing a cohort of 231 patients and a comprehensive gene pool of 12,571 genes, our models— XGBoost, ETC, and RFC—predicted the probability of surviving HCC beyond five years based on each patients’ genetic signatures. ETC achieved 95.3% accuracy, 92.1% precision, 97.2% recall, and an F1 score of 94.6%. RFC paralleled this with a 95.3% accuracy, 97.1% precision, 91.7% recall, and an F1 score of 94.3%, while XGBoost demonstrated 90.6% accuracy, precision at 100%, a recall of 77.78%, and an F1 score of 87.5%. The core discovery of this research lies in the explication of critical genes such as TOP3B, SSBP3, and COX7A2L through the lens of XAI. These genes have been identified as the most influential in HCC prognosis by each of the three employed AI models. SSBP3, in particular, was a consistent standout across all models, while COX7A2L was highlighted by both XGBoost and RFC, marking their importance in the context of HCC. Notably, despite XGBoost’s slightly lower F1 score, its interpretability through SHAP values is markedly superior, offering clearer insights into gene influence as evidenced by higher Shapley values (**Figure 3** A, B, and C). These pronounced explanatory points by XGBoost demonstrate its robust potential in deciphering complex biological interactions and contributing to the advancement of HCC diagnostics and therapeutics. Hence, the implementation of XAI in clinical settings necessitates a balanced consideration of the trade-offs between predictability and explainability offered by different XAI models. This delicate balance is pivotal in selecting the appropriate model for practical applications in the diagnosis and treatment of HCC, ensuring that the chosen model not only provides accurate predictions but also imparts meaningful insights for clinical decision-making.

The study’s XAI approach, leveraging Shapley values, provides a nuanced analysis of individual gene impacts on HCC patient prognosis, revealing key inflection points where gene influence becomes particularly pronounced, as detailed in **Figure 3** D, E, and F. These local explanations deliver precise targets for potential gene modulation, offering a pathway to significantly enhance patient survival rates. The findings serve as benchmarks for establishing normative biomarker levels within the body’s complex biological matrix. For example, the analysis delineates that TOP3 and COX7A2L expressions below approximately 3.5 and 8 transcripts per million respectively are associated with favorable HCC prognoses, while an SSBP3 expression above approximately 5.5 transcripts per million correlates with better survival outcomes. The detectability of these inflection points can vary based on the inherent explainability of the underlying data and AI algorithms. This study demonstrates variability in the detectability of these points, with the XGBoost model presenting a marked inflection for SSBP3 (**Figure 4** A), in contrast to a more nuanced indication from the ETC model (**Figure 4** B). This pattern of clarity versus nuance in the models’ local explanations is also evident in the analysis of other genes. The clarity of such local explanations, when aligned with overarching global insights, becomes critical in HCC where early detection is key yet challenging. This research underlines the necessity for a judicious balance between model predictability and explainability, particularly in clinical applications. The findings advocate that, notwithstanding the trade-offs, XAI models can revolutionize early diagnosis, accurate prognosis, and facilitate the advancement of tailored therapies, aligning treatment closely with the individual genetic profile of each patient’s tumor.

**Figure 4.**
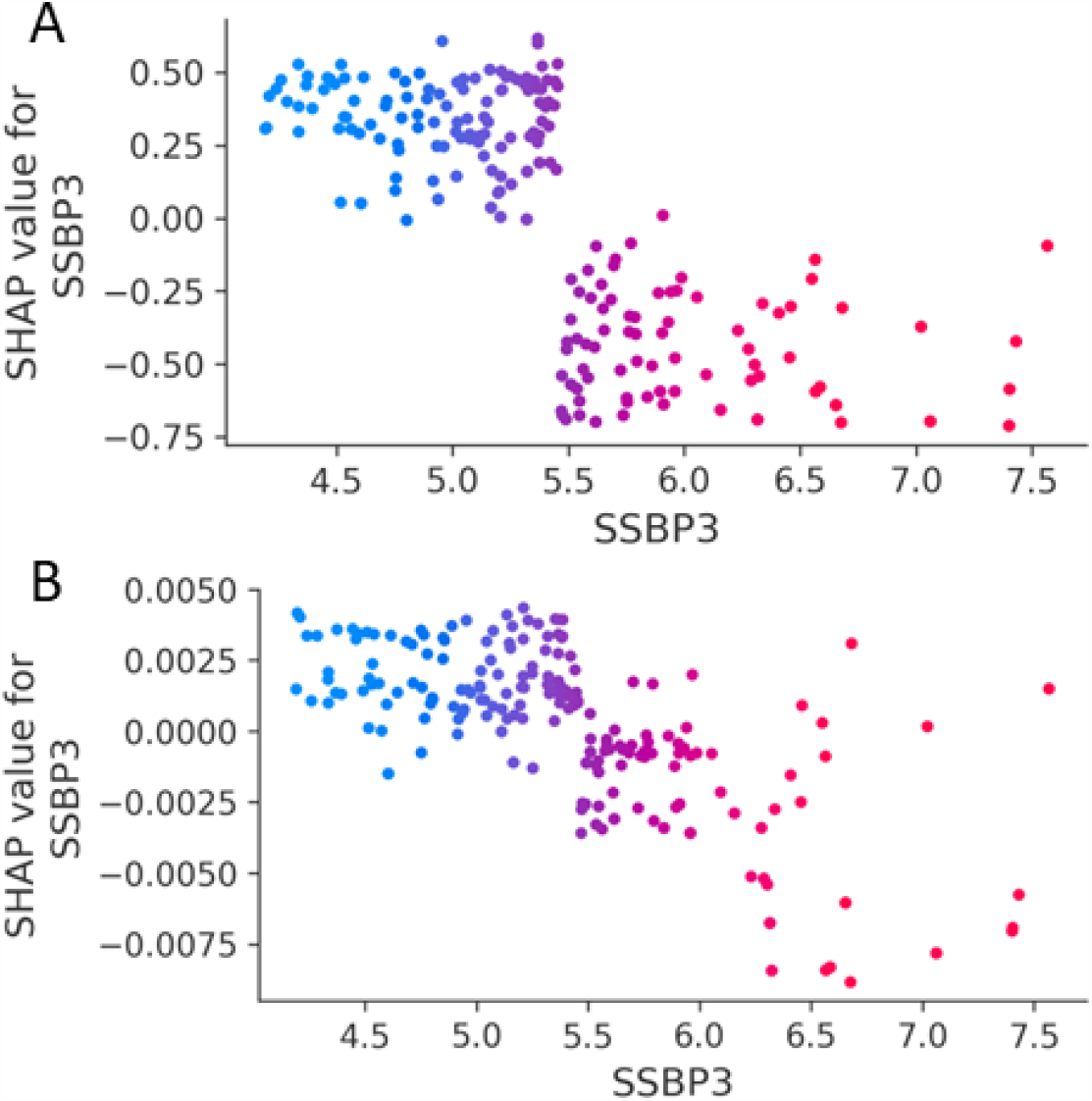
Comparison of local explanations between **A** XGBoost-SHAP and **B** ETC-SHAP for SSBP3 gene.

### Quantification of explainability

A significant obstacle in the application of XAI within medical contexts is the inherent speculative nature of the insights, which are often correlational rather than causational (*34, 35*). Acknowledging this challenge, our study introduces a probabilistic causal inference framework to quantify the impact of discoveries made by XAI, specifically relating to how gene features associate with patient outcomes in HCC. This methodology allows us to evaluate the potential for prognostic improvement by observing the fluctuations in gene expression levels as interpreted through XAI local explanations. By mathematically quantifying these relationships, we can gauge the probable efficacy of targeted therapeutic interventions, thereby enhancing the clinical relevance of XAI-derived insights. We formulate the probabilistic models to infer causality from the associative links revealed with XAI through the following equations:

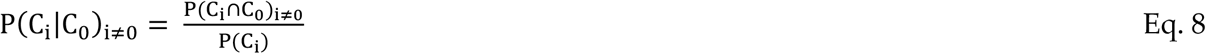

for individual cancer promoters/suppressors, and

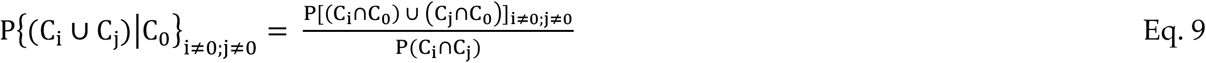

for combinatorial sets of cancer promoters/suppressors.

Where *C*_0_ is the number of patients in the cohort that survived beyond five years, *C*_*i*_ and *C*_*j*_ are the number of patients in the cohort that satisfy the genetic conditions (*i* and *j*) identified by XAI, and *P*(*C*_*i*_ ∩ *C*_*j*_) is the probability that conditions *i* and *j* are both satisfied.

Employing this probabilistic causal inference approach, our study quantifies the validity of hypotheses generated by XAI models in the context of HCC. Utilizing this framework, we can ascertain the likelihood of specific gene expression patterns being indicative of HCC prognosis. For instance, the probabilistic truth of the assertion that lower expressions of TOP3 and COX7A2L (below approximately 3.5 and 8 transcripts per million, respectively) are linked to better HCC outcomes, and higher SSBP3 expression (above approximately 5.5 transcripts per million) associates with improved survival, is computed to be 66.7%. This quantification not only reinforces the credibility of the insights derived from XAI but also substantiates the potential for these gene expressions to serve as predictive markers for patient prognosis.

### Comparative analysis of the predictability and explainability of genetic biomarkers identified by our XAI models against the established HCC biomarker, AFP

Predictability is evaluated using F1 scores, a harmonized measure of precision and recall, while explainability is appraised through local XAI explanations and the probabilistic causal inference framework discussed above. Our findings, detailed in **Table 1**, reveal that the individual predictability of TOP3B, SSBP3, and COX7A2L is robust, with scores of 77.6%, 80%, and 82.4%, respectively, juxtaposed with AFP’s predictability of 80.5% derived from XGBoost-SHAP based XAI models. Significantly, a composite biomarker panel incorporating TOP3B, SSBP3, and COX7A2L genes surpasses the predictive accuracy of AFP alone, achieving an 85.7% predictability rate. This enhancement suggests the potential superiority of a multi-biomarker approach over the traditional singular use of AFP. Conversely, the explainability of AFP is unquantifiable due to the absence of discernible inflection points in the SHAP local explanation plot (**Figure 5**). In contrast, the combined explainability of TOP3B, SSBP3, and COX7A2L, as reflected by our XAI’s inflection points, is quantified at 66.7%, advocating for their integration into a more comprehensive diagnostic and therapeutic strategy for HCC.

**Table 1.**
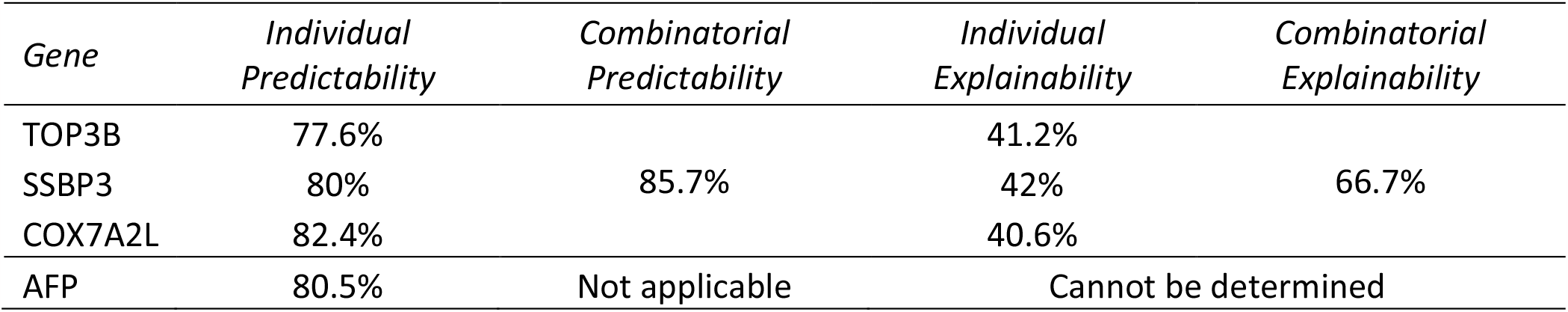
Summary of individual and combinatorial predictability and explainability of the top genetic biomarkers identified by our XAI models against the established HCC biomarker, AFP.

| *Gene* | *Individual Predictability* | *Combinatorial Predictability* | *Individual Explainability* | *Combinatorial Explainability* |
| --- | --- | --- | --- | --- |
| TOP3B | 77.6% | 85.7% | 41.2% | 66.7% |
| SSBP3 | 80% |  | 42% |  |
| COX7A2L | 82.4% |  | 40.6% |  |
| AFP | 80.5% | Not applicable | Cannot be determined |  |

**Figure 5.**
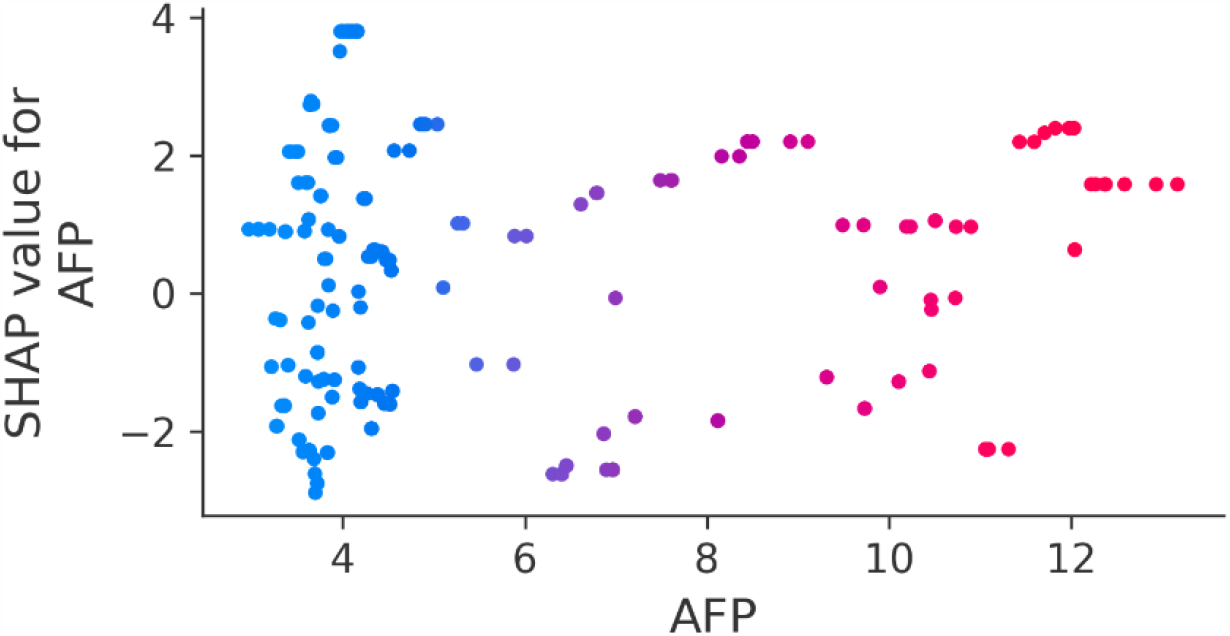
Local explanations with XGBoost-SHAP for AFP gene revealing the absence of discernible inflection points.

### Traditional Analysis of XAI generated biomarkers with GEPIA2

The use of advanced XAI methods in HCC diagnostic and therapeutic research has highlighted certain biomarkers (TOP3B, SSBP3, and COX7A2L) as highly influential. Upon evaluation using established survival analysis methods, including Kaplan-Meier plots and pertinent statistical tests (log-rank test, Mantel–Cox test, and Cox proportional hazards model, with a 95% confidence interval), two of the three identified biomarkers, TOP3B and COX7A2L, demonstrate statistically significant prognostic value in HCC. This assessment was conducted in GEPIA2 (*36*), which comprises a dataset of 50 normal liver tissue samples and a total of 419 liver samples, using custom cutoff values based on XAI results (**Figure 6**). The absence of statistical significance is observed not only in SSBP3 but also in AFP, a well-established biomarker for HCC. This divergence in results obtained from XAI methods compared to those from traditional survival analysis underscores the complexities and potential challenges inherent in interpreting genomic data.

**Figure 6.**
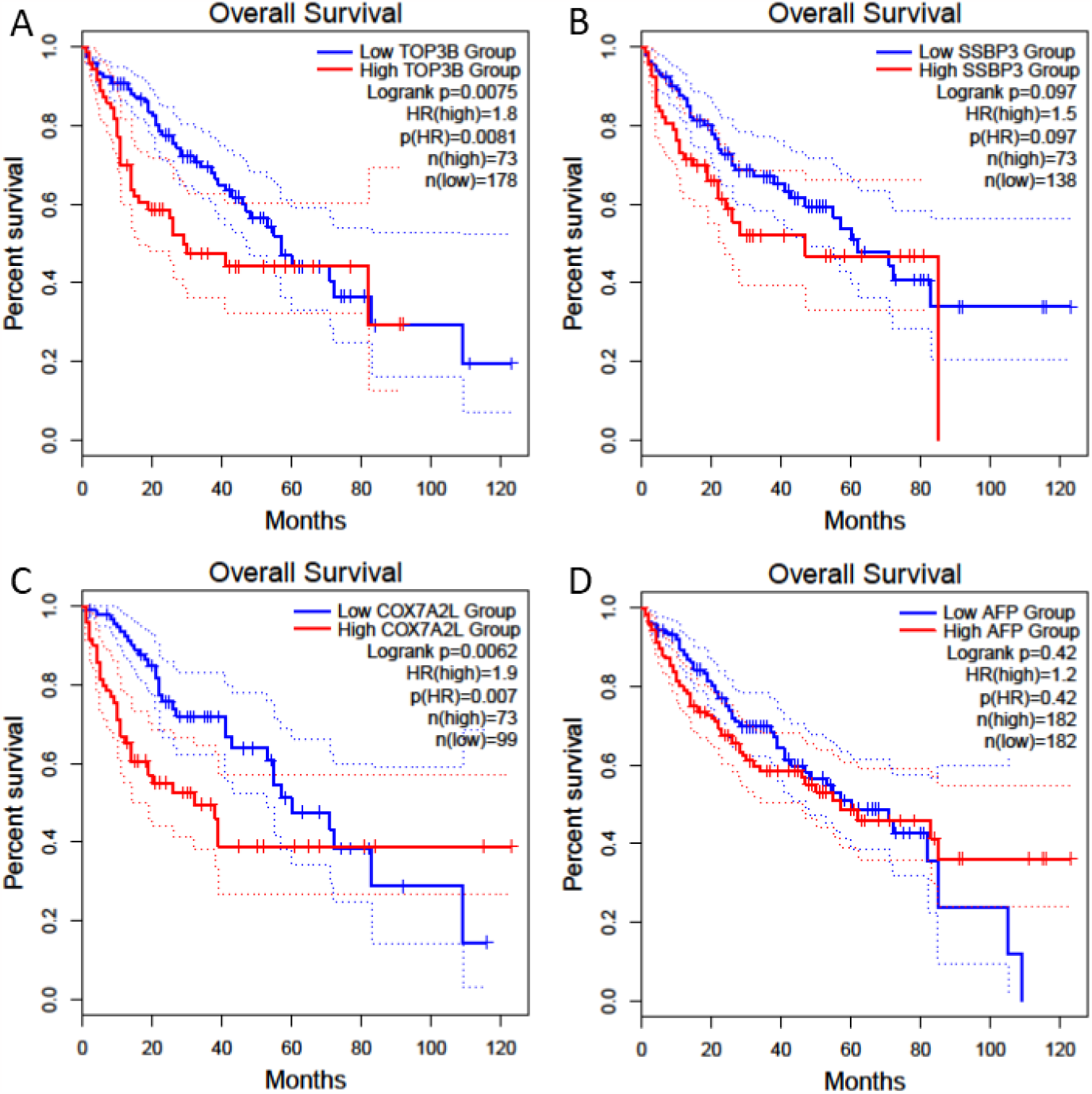
Survival analysis was conducted on biomarkers identified as influential by XAI, namely **A** TOP3B, **B** SSBP3, **C** COX7A2L, and the established HCC biomarker, **D** AFP, using the GEPIA2 database. The group cutoff values were set at 80% and 49% for TOP3B, 80% and 38% for SSBP3, 80% and 27% for COX7A2L, while AFP utilized a balanced 50% cutoff. The median cutoff was chosen for AFP as the XAI-generated local explanations did not exhibit any clear inflection points, in contrast to the patterns observed for TOP3B, SSBP3, and COX7A2L.

While XAI can offer insights into complex data patterns and feature importance, traditional statistical methods anchor these findings within established scientific protocols that may not capture the multifaceted nature of genomic influences on disease outcomes. The minimal variance in results yielded by these two analytical methods underscores the necessity of employing a multifaceted approach to comprehensively evaluate the prognostic significance of genetic markers in intricate diseases such as HCC. This approach highlights the importance of integrating diverse methodologies for a more thorough understanding and assessment. This integrative approach may lead to a more robust understanding and better clinical decision-making by accounting for the strengths and limitations of each method.

Based on our analysis, it appears that traditional statistical methods for identifying influential biomarkers in HCC may treat the problem as linear and one-dimensional. This approach might be insufficient for capturing the complexity inherent in the biological interactions that underlie HCC. In reality, the identification of critical biomarkers is a non-linear and multi-dimensional challenge that requires the nuanced analysis provided by advanced computational techniques. XAI methods offer a powerful alternative by accommodating the non-linear and complex nature of genomic data, evident from the local explanations in **Figure 3**. XAI not only facilitates the discovery of intricate patterns within high-dimensional datasets but also provides insight into the reasoning behind these findings, allowing for a better understanding of each biomarker’s role in HCC prognosis. Therefore, our findings suggest a significant advantage in using XAI over traditional methods for prognostic evaluations in HCC. This also underscores the importance of adopting more sophisticated analytical tools in genomic research to improve the accuracy and interpretability of results, which could lead to enhanced clinical decision-making and patient outcomes.

### Relevance of the XAI-discovered influential biomarkers to the Hispanic demographic

The pertinence of the novel prognostic genes for the Hispanic demographic is further emphasized by a supplementary analysis of a localized dataset (*27*). This analysis seeks to highlight the characteristics of these genes within the Hispanic population (*n=13*) contrasted against a TCGA-based non-Hispanic cohort (*n=50*). As illustrated in **Figure 7**, a marked statistical significance in the expression levels of TOP3B and COX7A2L between non-tumor and tumor samples has been observed (p < 0.05) in both the Hispanic and TCGA-based non-Hispanic cohorts. This significant differential expression suggests a potential role for these genes in the pathophysiology of HCC. These findings may indicate that TOP3B and COX7A2L have the potential to act as biomarkers or therapeutic targets for HCC in both population groups. Furthermore, these results are in alignment with our XAI-based discoveries that pertain to the general population. Such alignment reinforces the validity of these genes as critical elements in the understanding and treatment of HCC.

**Figure 7.**
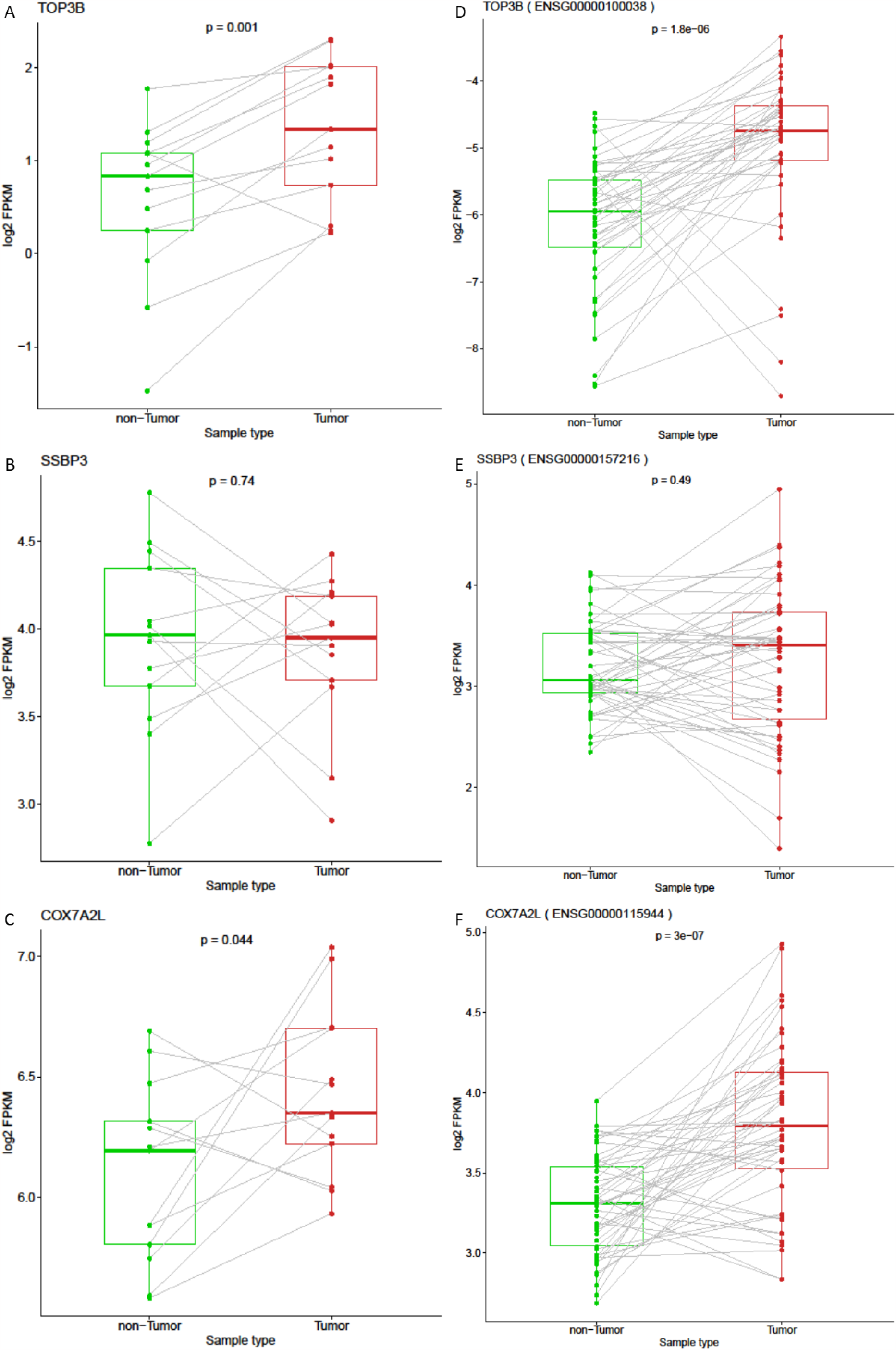
Box plot depicting the comparative expression levels of **A**. TOP3, **B**. SSBP3, and **C**. COX7A2L in non-tumor versus tumor HCC samples within the Hispanic population, contrasted against the expression profiles of **D**. TOP3, **E**. SSBP3, and **F**. COX7A2L in non-tumor versus tumor HCC samples from the TCGA-based non-Hispanic cohort.

## 4. Discussion

The application of XAI in this study has significantly advanced the predictive modeling of five-year survivability rates in HCC patients by utilizing gene expression profiles. Through XAI, we have pinpointed novel genes with considerable prognostic significance, with TOP3B, SSBP3, and COX7A2L emerging as key biomarkers. These biomarkers have demonstrated superiority for HCC diagnosis and prognosis than the traditionally relied-upon biomarker, AFP. The influence of these biomarkers is meticulously documented in the previous section and is contextualized below within the expansive sphere of HCC research, highlighting their potential in refining the accuracy of HCC diagnostics and enriching the prognostic landscape.

**DNA Topoisomerase III Beta (TOP3B)** is a versatile DNA topoisomerase that operates on both RNA and DNA and is localized within the nucleus and cytoplasm (*37*–*39*). While localized in the nucleus, TOP3B associates with Tudor Domain–Containing 3 Protein (TDRD3) to alleviate negative supercoiling of double stranded DNA, as well as settling R-loops (*39*), thus facilitating efficient transcription. TOP3B catalyzes the transient separation and connection of a single strand of DNA thus relaxing the supercoils and changing the topology of DNA by permitting the strands to pass by one another (*37, 39*). The enzyme also interacts with SGS1, a DNA helicase, further partaking in DNA recombination, genome stability, and cell aging (*38, 39*). Within the cytoplasm, the TOP3B-TDRD3 complex works to monitor mRNA topological stress and translation (*39*). Notably, diminished TOP3B expression has been correlated with improved survival outcomes in breast cancer, suggesting its potential as a prognostic biomarker in oncology (*38*).

**Single Stranded DNA Binding Protein 3 (SSBP3)** belongs to a family of single stranded DNA binding proteins that enable pyrimidine-rich single-stranded DNA binding and transcription coactivator activity (*40*–*42*). Out of the SSBP3 gene there are three transcripts in the human: SSBP3a, SSBP3b, SSBP3c; and downregulation of SSBP3 shows a decrease in cell proliferation that causes neuroectoderm tissue loss (*41*). It has also been linked to diabetes as SSBP3 directs the proper function of pancreatic islets. A decrease or loss of β-cell SSBP3 causes changes to the β-cell function and quantity (*42*). SSBP3 transcription co-regulators direct proper islet function needed for glucose homeostasis (*42*). Diabetes mellitus (DM) is a risk factor of HCC and has been linked to advanced lesions in patients with HCC (*43*). Chromosomal localization by SSBP3 and the putative single-stranded DNA-binding activity suggest that all three members of this family are capable of tumor suppressor activity by gene dosage or other epigenetic mechanisms (*44*).

**Cytochrome C Oxidase Subunit 7A2 Like (COX7A2L)** is a nuclear DNA-encoded gene involved in the assembly of mitochondrial respiratory chain through binding complex III that stabilizes the III2+IV supercomplex (*45*). Increased COX7A2L expression in the muscle has been associated with lower body fat and improved cardiorespiratory fitness in humans (*46*). Interestingly, such overexpression was also shown to promote tumor growth and metastasis by inducing ROS production in HCC cells (*47*).

To summarize, the literature suggests that TOP3B, involved in DNA and RNA topological changes, is tied to improved survival in breast cancer, hinting at its broader oncological relevance. SSBP3, which assists in single-stranded DNA binding and is crucial for cell proliferation and pancreatic function, also exhibits potential tumor suppressor activities. Its downregulation is linked to diabetes, a known HCC risk factor. COX7A2L, related to mitochondrial function and associated with physical fitness, paradoxically may encourage HCC progression through reactive oxygen species production. The discovery of TOP3B, SSBP3, and COX7A2L as novel prognostic biomarkers using XAI aligns with existing research and enriches the current knowledge base concerning HCC’s underlying mechanisms. By leveraging gene expression profiles, XAI has not only highlighted these genes’ predictive value for five-year survivability rates but also emphasized their potential to surpass AFP in diagnostic and prognostic accuracy. Such integration of cutting-edge technology with genomic data promises to refine current approaches to HCC management, potentially leading to earlier detection, more personalized treatment plans, and better overall patient outcomes.

## 5. Conclusions

This study computationally explored the relationships between 12571 genes and corresponding expressions with the survival duration of 231 patients. Central to the findings of this study, the XAI analysis highlighted a set of pivotal genes including TOP3B, SSBP3, and COX7A2L. Notably, SSBP3 emerged as a key gene across all three AI models, and COX7A2L was singled out by two out of three AI models, underscoring their potential significance in HCC prognosis. For future research or clinical practice, we strongly recommend the use of multiple biomarkers in tandem for HCC diagnosis and treatment that we discovered using a XAI model framework. Combining multiple biomarkers for HCC diagnosis and treatment is crucial for: improved accuracy, early detection, confirmation of diagnosis, prognosis and staging, monitoring treatment response, detection of recurrence, personalized medicine, and most importantly reduction of false positives/negatives.

The exigency for improved HCC diagnostics is paramount, given the limitations of current methods such as ultrasound and solitary AFP testing. This study advocates for a panel of combinatorial biomarkers, inferenced through XAI, to enhance the precision of HCC care continuum—from initial screening to treatment and post-therapy monitoring. While these combinatorial biomarkers present a promising leap beyond the traditional techniques, necessitating the transition from ultrasound and standalone AFP screenings, they require rigorous validation. The path to clinical adoption is fraught with logistical challenges that must be navigated to unlock the full potential of this approach for transforming HCC patient management and outcomes.

This study also underscores the necessity of employing advanced computational methods to address the multi-dimensional challenges of genomic data interpretation and the advantage of XAI in uncovering and understanding the prognostic value of novel biomarkers in HCC. The results from XAI models not only outperform the predictive capabilities of traditional biomarkers such as AFP but also provide deeper insights into the gene expression profiles that could serve as potential targets for early detection and personalized therapy. The effectiveness of tree-based ensemble AI models in predicting patient outcomes was validated against traditional statistical tests, which often fail to capture the complex interrelations of gene expressions and prognostic factors. The contrast between XAI predictions and conventional methods like Kaplan-Meier plots and log-rank tests further emphasizes the multifaceted nature of gene influence on disease progression, suggesting that a multi-dimensional approach to biomarker discovery is necessary for a more accurate and comprehensive understanding of HCC.

From a broader biological perspective, by harnessing XAI, researchers can gain a more profound understanding of genetic datasets, potentially guiding the development of targeted therapies that could precisely modulate gene expression to improve patient prognosis. The results emphasize the need for further exploration of the roles of the influential biomarkers for HCC (identified by XAI) and their interactions with the immune system and tumor cells, which may open new avenues for cancer treatment.

In a bid to catalyze further research and hasten the clinical integration of our findings, we have made the dataset on which our XAI models were trained publicly available. Additionally, we provide access to the refined XAI models that incorporate the identified influential genes. These models are ready for application on new data from diverse cohorts, offering a valuable tool for prognostic assessment in HCC. This initiative is aimed at fostering collaborative validation and adaptation across the scientific community, thereby advancing the precision and personalization of HCC treatment strategies. *Note to editor(s) and reviewer(s): The authors pledge that the data, code, and trained models will be made available on Code Ocean platform once accepted for publication (link will be provided here before the final version is published by the journal)*.

## 6. Limitations

Serum AFP levels are commonly used for HCC diagnosis. Owing to the absence of serum AFP data for our selected cohorts, we have resorted to measuring AFP mRNA levels instead. This approach is substantiated by existing literature that recognizes AFP mRNA as a diagnostic and prognostic marker (*48*–*50*). It is vital to note that although we have utilized a cutting-edge multi-model XAI, a probabilistic causal inference framework, and various data sources to identify and validate new prognostic biomarkers for HCC, our conclusions grounded in bioinformatics may still need additional validation through wet-lab experiments prior to their integration into mainstream clinical practices and therapeutic strategies.

## Abbreviations

HCC: Hepatocellular carcinoma
AI: Artificial Intelligence
XAI: eXplainable Artificial Intelligence
XGBoost: eXtreme Gradient Boosting
RFC: Random Forest Classifier
ETC: Extra Trees Classifier
SHAP: SHapley Additive exPlanations
TOP3B: DNA Topoisomerase III Beta
SSBP3: Single-stranded DNA binding protein 3
COX7A2L: Cytochrome C Oxidase Subunit 7A2 Like
AFP: Alpha-fetoprotein

## Authors’ contributions

Conceptualization – EGC, YB, DC; Analysis – All authors; Writing – EGC, DC, YB.

## Data Availability

All relevant data, code, and trained models will be made available on Code Ocean platform once accepted for publication (link will be provided by the authors before the final version is published by the journal).

## Acknowledgement

The RNA-seq data derived from liver samples of South Texas Hispanic patients with HCC were obtained by Dr. LuZhe Sun and his co-workers with the support of a grant from Clayton Foundation for Research to Drs. Francisco Cigarroa and LuZhe Sun. We also extend our sincere gratitude to Dr. Robin Leach for her insightful discussions on the landscape of HCC biomarkers and to Dr. Hakan Başağaoğlu for his critical review of our study.

## References

1. F. Bray, J. Ferlay, I. Soerjomataram, R. L. Siegel, L. A. Torre, A. Jemal, Global cancer statistics 2018: GLOBOCAN estimates of incidence and mortality worldwide for 36 cancers in 185 countries. CA A Cancer J Clinicians 68, 394–424 (2018).

2. K. D. Miller, A. Goding Sauer, A. P. Ortiz, S. A. Fedewa, P. S. Pinheiro, G. Tortolero-Luna, D. Martinez-Tyson, A. Jemal, R. L. Siegel, Cancer Statistics for Hispanics/Latinos, 2018. CA A Cancer J Clinicians 68, 425–445 (2018).

3. R. L. Siegel, K. D. Miller, N. S. Wagle, A. Jemal, Cancer statistics, 2023. CA A Cancer J Clinicians 73, 17–48 (2023).

4. M. Kudo, G. Han, R. S. Finn, R. T. P. Poon, J.-F. Blanc, L. Yan, J. Yang, L. Lu, W.-Y. Tak, X. Yu, J.-H. Lee, S.-M. Lin, C. Wu, T. Tanwandee, G. Shao, I. B. Walters, C. Dela Cruz, V. Poulart, J.-H. Wang, Brivanib as adjuvant therapy to transarterial chemoembolization in patients with hepatocellular carcinoma: A randomized phase III trial: HEPATOLOGY, Vol. XX, No. X, 2014 KUDO ET AL. Hepatology 60, 1697–1707 (2014).

5. M. Chen, B. Zhang, W. Topatana, J. Cao, H. Zhu, S. Juengpanich, Q. Mao, H. Yu, X. Cai, Classification and mutation prediction based on histopathology H&E images in liver cancer using deep learning. npj Precis. Onc. 4, 14 (2020).

6. Q. Li, J. Wu, M. Zhu, Y. Tang, L. Jin, Y. Chen, M. Jin, Z. Peng, A novel risk signature based on autophagy-related genes to evaluate tumor immune microenvironment and predict prognosis in hepatocellular carcinoma. Computers in Biology and Medicine 152, 106437 (2023).

7. N. D. Parikh, N. Tayob, A. G. Singal, Blood-based biomarkers for hepatocellular carcinoma screening: Approaching the end of the ultrasound era? Journal of Hepatology 78, 207–216 (2023).

8. A. Walakira, C. Skubic, N. Nadižar, D. Rozman, T. Režen, M. Mraz, M. Moškon, Integrative computational modeling to unravel novel potential biomarkers in hepatocellular carcinoma. Computers in Biology and Medicine 159, 106957 (2023).

9. P. R. Galle, A. Forner, J. M. Llovet, V. Mazzaferro, F. Piscaglia, J.-L. Raoul, P. Schirmacher, V. Vilgrain, EASL Clinical Practice Guidelines: Management of hepatocellular carcinoma. Journal of Hepatology 69, 182–236 (2018).

10. P. R. Galle, F. Foerster, M. Kudo, S. L. Chan, J. M. Llovet, S. Qin, W. R. Schelman, S. Chintharlapalli, P. B. Abada, M. Sherman, A. X. Zhu, Biology and significance of alpha-fetoprotein in hepatocellular carcinoma. Liver International 39, 2214–2229 (2019).

11. F. Özdemir, A. Baskiran, The Importance of AFP in Liver Transplantation for HCC. J Gastrointest Canc 51, 1127–1132 (2020).

12. Z. Wei, Y. Zhang, H. Lu, J. Ying, H. Zhao, J. Cai, Serum alpha-fetoprotein as a predictive biomarker for tissue alpha-fetoprotein status and prognosis in patients with hepatocellular carcinoma. Transl Cancer Res 11, 669–677 (2022).

13. C. Saitta, G. Raffa, A. Alibrandi, S. Brancatelli, D. Lombardo, G. Tripodi, G. Raimondo, T. Pollicino, PIVKA-II is a useful tool for diagnostic characterization of ultrasound-detected liver nodules in cirrhotic patients. Medicine 96, e7266 (2017).

14. P. Luo, S. Wu, Y. Yu, X. Ming, S. Li, X. Zuo, J. Tu, Current Status and Perspective Biomarkers in AFP Negative HCC: Towards Screening for and Diagnosing Hepatocellular Carcinoma at an Earlier Stage. Pathol. Oncol. Res. 26, 599–603 (2020).

15. G. Zhang, S.-A. Ha, H. K. Kim, J. Yoo, S. Kim, Y. S. Lee, S. Y. Hur, Y. W. Kim, T. E. Kim, Y. G. Park, J. Wang, Y. Yang, Z. Xu, E. Y. Song, Z. Huang, P. Jirun, J. Zhongtian, Q. Shishi, C. Zhuqingqing, G. Lei, J. W. Kim, Combined Analysis of AFP and HCCR-1 as an Useful Serological Marker for Small Hepatocellular Carcinoma: A Prospective Cohort Study. Disease Markers 32, 265–271 (2012).

16. F. De Stefano, E. Chacon, L. Turcios, F. Marti, R. Gedaly, Novel biomarkers in hepatocellular carcinoma. Digestive and Liver Disease 50, 1115–1123 (2018).

17. M. Sherman, Current status of alpha-fetoprotein testing. Gastroenterol Hepatol (N Y) 7, 113–114 (2011).

18. K. Tzartzeva, J. Obi, N. E. Rich, N. D. Parikh, J. A. Marrero, A. Yopp, A. K. Waljee, A. G. Singal, Surveillance Imaging and Alpha Fetoprotein for Early Detection of Hepatocellular Carcinoma in Patients With Cirrhosis: A Meta-analysis. Gastroenterology 154, 1706–1718.e1 (2018).

19. K. Zhu, Z. Dai, J. Zhou, Biomarkers for hepatocellular carcinoma: progression in early diagnosis, prognosis, and personalized therapy. Biomark Res 1, 10 (2013).

20. L. Huang, Z. Songyang, Z. Dai, Y. Xiong, Field cancerization profile-based prognosis signatures lead to more robust risk evaluation in hepatocellular carcinoma. iScience 25, 103747 (2022).

21. H. Sung, J. Ferlay, R. L. Siegel, M. Laversanne, I. Soerjomataram, A. Jemal, F. Bray, Global Cancer Statistics 2020: GLOBOCAN Estimates of Incidence and Mortality Worldwide for 36 Cancers in 185 Countries. CA A Cancer J Clin 71, 209–249 (2021).

22. P. Johnson, Q. Zhou, D. Y. Dao, Y. M. D. Lo, Circulating biomarkers in the diagnosis and management of hepatocellular carcinoma. Nat Rev Gastroenterol Hepatol 19, 670–681 (2022).

23. M.-C. Jiang, J.-J. Ni, W.-Y. Cui, B.-Y. Wang, W. Zhuo, Emerging roles of lncRNA in cancer and therapeutic opportunities. Am J Cancer Res 9, 1354–1366 (2019).

24. J. M. Llovet, R. K. Kelley, A. Villanueva, A. G. Singal, E. Pikarsky, S. Roayaie, R. Lencioni, K. Koike, J. Zucman-Rossi, R. S. Finn, Hepatocellular carcinoma. Nat Rev Dis Primers 7, 6 (2021).

25. Y. N. Flores, G. D. Datta, L. Yang, E. Corona, D. Devineni, B. A. Glenn, R. Bastani, F. P. May, Disparities in Hepatocellular Carcinoma Incidence, Stage, and Survival: A Large Population-Based Study. Cancer Epidemiology, Biomarkers & Prevention 30, 1193–1199 (2021).

26. V. Kulasingam, E. P. Diamandis, Strategies for discovering novel cancer biomarkers through utilization of emerging technologies. Nat Rev Clin Oncol 5, 588–599 (2008).

27. G. Zheng, H. Bouamar, M. Cserhati, C. R. Zeballos, I. Mehta, H. Zare, L. Broome, R. Hu, Z. Lai, Y. Chen, F. E. Sharkey, M. Rani, G. A. Halff, F. G. Cigarroa, L. Sun, Integrin alpha 6 is upregulated and drives hepatocellular carcinoma progression through integrin α6β4 complex. Intl Journal of Cancer 151, 930–943 (2022).

28. Q. Lian, S. Wang, G. Zhang, D. Wang, G. Luo, J. Tang, L. Chen, J. Gu, HCCDB: A Database of Hepatocellular Carcinoma Expression Atlas. Genomics, Proteomics & Bioinformatics 16, 269–275 (2018).

29. T. Chen, C. Guestrin, “XGBoost: A Scalable Tree Boosting System” in Proceedings of the 22nd ACM SIGKDD International Conference on Knowledge Discovery and Data Mining (ACM, San Francisco California USA, 2016; https://dl.acm.org/doi/10.1145/2939672.2939785), pp. 785–794.

30. S. M. Lundberg, G. Erion, H. Chen, A. DeGrave, J. M. Prutkin, B. Nair, R. Katz, J. Himmelfarb, N. Bansal, S.-I. Lee, From local explanations to global understanding with explainable AI for trees. Nat Mach Intell 2, 56–67 (2020).

31. D. Chakraborty, C. Ivan, P. Amero, M. Khan, C. Rodriguez-Aguayo, H. Başağaoğlu, G. Lopez-Berestein, Explainable Artificial Intelligence Reveals Novel Insight into Tumor Microenvironment Conditions Linked with Better Prognosis in Patients with Breast Cancer. Cancers (Basel) 13, 3450 (2021).

32. J. Meena, Y. Hasija, Application of explainable artificial intelligence in the identification of Squamous Cell Carcinoma biomarkers. Computers in Biology and Medicine 146, 105505 (2022).

33. S. Rajpal, A. Rajpal, A. Saggar, A. K. Vaid, V. Kumar, M. Agarwal, N. Kumar, XAI-MethylMarker: Explainable AI approach for biomarker discovery for breast cancer subtype classification using methylation data. Expert Systems with Applications 225, 120130 (2023).

34. J. G. Richens, C. M. Lee, S. Johri, Improving the accuracy of medical diagnosis with causal machine learning. Nat Commun 11, 3923 (2020).

35. M. Prosperi, Y. Guo, M. Sperrin, J. S. Koopman, J. S. Min, X. He, S. Rich, M. Wang, I. E. Buchan, J. Bian, Causal inference and counterfactual prediction in machine learning for actionable healthcare. Nat Mach Intell 2, 369–375 (2020).

36. Z. Tang, B. Kang, C. Li, T. Chen, Z. Zhang, GEPIA2: an enhanced web server for large-scale expression profiling and interactive analysis. Nucleic Acids Res 47, W556–W560 (2019).

37. Y. Pommier, A. Nussenzweig, S. Takeda, C. Austin, Human topoisomerases and their roles in genome stability and organization. Nat Rev Mol Cell Biol 23, 407–427 (2022).

38. F. Moreira, M. Arenas, A. Videira, F. Pereira, Molecular Evolution of DNA Topoisomerase III Beta (TOP3B) in Metazoa. J Mol Evol 89, 384–395 (2021).

39. Y. Bai, L.-D. Li, J. Li, X. Lu, Targeting of topoisomerases for prognosis and drug resistance in ovarian cancer. J Ovarian Res 9, 35 (2016).

40. C. Maffeo, A. Aksimentiev, Molecular mechanism of DNA association with single-stranded DNA binding protein. Nucleic Acids Research 45, 12125–12139 (2017).

41. Z. Yin, K. Zhang, X. Peng, Z. Jiang, W. Yuan, Y. Wang, Y. Li, X. Ye, Y. Dong, Y. Wan, B. Ni, P. Zhu, X. Fan, X. Wu, X. Mo, SIVA1 Regulates the Stability of Single-Stranded DNA-Binding Protein 3 Isoforms. Mol Biol 52, 707–714 (2018).

42. E. Toren, J. D. Kepple, K. V. Coutinho, S. O. Poole, I. M. Deeba, T. H. Pierre, Y. Liu, M. M. Bethea, C. S. Hunter, The SSBP3 co-regulator is required for glucose homeostasis, pancreatic islet architecture, and beta-cell identity. Molecular Metabolism 76, 101785 (2023).

43. D. N. Amarapurkar, N. D. Patel, P. M. Kamani, Impact of diabetes mellitus on outcome of HCC. Annals of Hepatology 7, 148–151 (2008).

44. P. Castro, H. Liang, J. C. Liang, L. Nagarajan, A Novel, Evolutionarily Conserved Gene Family with Putative Sequence-Specific Single-Stranded DNA-Binding Activity. Genomics 80, 78–85 (2002).

45. R. Pérez-Pérez, T. Lobo-Jarne, D. Milenkovic, A. Mourier, A. Bratic, A. García-Bartolomé, E. Fernández-Vizarra, S. Cadenas, A. Delmiro, I. García-Consuegra, J. Arenas, M. A. Martín, N.-G. Larsson, C. Ugalde, COX7A2L Is a Mitochondrial Complex III Binding Protein that Stabilizes the III2+IV Supercomplex without Affecting Respirasome Formation. Cell Reports 16, 2387–2398 (2016).

46. G. Benegiamo, M. Bou Sleiman, M. Wohlwend, S. Rodríguez-López, L. J. E. Goeminne, P.-P. Laurila, M. Klevjer, M. K. Salonen, J. Lahti, P. Jha, S. Cogliati, J. A. Enriquez, B. M. Brumpton, A. Bye, J. G. Eriksson, J. Auwerx, COX7A2L genetic variants determine cardiorespiratory fitness in mice and human. Nat Metab 4, 1336–1351 (2022).

47. G. Wang, B. Popovic, J. Tao, A. Jiang, Overexpression of COX7RP promotes tumor growth and metastasis by inducing ROS production in hepatocellular carcinoma cells. Am J Cancer Res 10, 1366–1383 (2020).

48. M. Matsumura, Y. Shiratori, Y. Niwa, T. Tanaka, K. Ogura, T. Okudaira, M. Imamura, K. Okano, S. Shiina, M. Omata, Presence of α-fetoprotein mRNA in blood correlates with outcome in patients with hepatocellular carcinoma. Journal of Hepatology 31, 332–339 (1999).

49. I. Wong, Quantitative comparison of alpha-fetoprotein and albumin mRNA levels in hepatocellular carcinoma/adenoma, non-tumor liver and blood: implications in cancer detection and monitoring. Cancer Letters 156, 141–149 (2000).

50. E. N. Debruyne, J. R. Delanghe, Diagnosing and monitoring hepatocellular carcinoma with alpha-fetoprotein: new aspects and applications. Clin Chim Acta 395, 19–26 (2008).

